# A deficiency screen of the *X* chromosome for Rap1 GTPase dominant interacting genes in *Drosophila* border cell migration

**DOI:** 10.1101/2025.01.02.631095

**Authors:** C. Luke Messer, Emily Burghardt, Jocelyn A. McDonald

**Affiliations:** Division of Biology, Kansas State University, Manhattan, KS 66506, USA; The University of Virginia’s College at Wise, Wise, VA 24293, USA

**Keywords:** Drosophila, border cells, collective cell migration, Rap1 GTPase, frizzled 4, Ubiquitin specific protease 16/45, strawberry notch

## Abstract

Collective cell migration is critical to embryonic development, wound healing, and the immune response, but also drives tumor dissemination. Understanding how cell collectives coordinate migration in vivo has been a challenge, with potential therapeutic benefits that range from addressing developmental defects to designing targeted cancer treatments. The small GTPase Rap1 has emerged as a regulator of both embryogenesis and cancer cell migration. How active Rap1 coordinates downstream signaling functions required for coordinated collective migration is poorly understood. *Drosophila* border cells undergo a stereotyped and genetically tractable in vivo migration within the developing egg chamber of the ovary. This group of 6-8 cells migrates through a densely packed tissue microenvironment and serves as an excellent model for collective cell migration during development and disease. Rap1, like all small GTPases, has distinct activity state switches that link extracellular signals to organized cell behaviors. Proper regulation of Rap1 activity is essential for successful border cell migration yet the signaling partners and other downstream effectors are poorly characterized. Using the known requirement for Rap1 in border cell migration, we conducted a dominant suppressor screen for genes whose heterozygous loss modifies the migration defects observed upon constitutively active *Rap1^V12^* expression. Here we identified seven genomic regions on the *X* chromosome that interact with *Rap1^V12^*. We mapped three genomic regions to single Rap1-interacting genes, *frizzled 4*, *Ubiquitin specific protease 16/45*, and *strawberry notch*. Thus, this unbiased screening approach identified multiple new candidate regulators of Rap1 activity with roles in collective border cell migration.

## Introduction

The small GTPase Rap1 has important roles in tissue morphogenesis and integrity, single cell and collective cell migration, wound healing, and tumor invasion in cancer (Jaśkiewicz et al., 2018; Kim et al., 2022; Messer and McDonald, 2023; Perez-Vale et al., 2022; Rothenberg et al., 2023; Sawant et al., 2018; Ueda et al., 2023; Volovetz et al., 2020; Zhang et al., 2017). Rap1, like other small GTPases, acts as a molecular switch with discrete “on” and “off” states. GTPase activity is regulated by a combination of GTPase activating proteins (GAPs) that speed up GTP hydrolysis and result in inactive GDP-bound GTPases, and guanine nucleotide exchange factors (GEFs) that promote dissociation of GDP allowing GTP to bind (Cherfils and Zeghouf, 2013; Raaijmakers and Bos, 2009; Zegers and Friedl, 2014). Several key regulators have been identified for Rap1, including the GAP Rapgap1 and the GEF PDZ-GEF/Dizzy (Boettner and Van Aelst, 2009; Jaśkiewicz et al., 2018; Sawant et al., 2018; Wang et al., 2013). Other signaling partners downstream of Rap1, including Afadin/Canoe and Rap1 interacting adaptor molecule (RIAM), promote Rap1-dependent cellular processes such as cell polarity and adhesion (Bonello et al., 2018; Bromberger et al., 2021; Hiremath et al., 2023; Rothenberg et al., 2023; Su et al., 2015; Walther et al., 2018). Despite progress in identifying some Rap1-effectors, our understanding of how Rap1 coordinates a diverse set of functions in a wide range of tissues remains limited, with many Rap1 roles unaccounted for by these known effectors.

To address this gap, we performed an unbiased dominant genetic interaction screen to identify downstream targets of Rap1 and other Rap1-interacting genes in a migrating cell collective, the *Drosophila* border cells. Similar approaches in *Drosophila* models of collective migration and morphogenesis have underscored the power of this technique to quickly identify candidate interacting genes that may otherwise be difficult to uncover (Chang et al., 2018; Geisbrecht et al., 2013; Hurd et al., 2013; McDonald et al., 2003; Patch et al., 2009; Ward et al., 2003). Specifically, we took advantage of the established role of Rap1 in regulating border cell migration, a genetically tractable in vivo model of collective cell migration (Chang et al., 2018; Sawant et al., 2018). Border cells are a group of 6-8 epithelial cells that are specified and recruited as a migratory cohort (cluster) during stages 8-9 of oogenesis. Subsequently, the border cell cluster delaminates from the epithelium, then migrates between the germline derived nurse cells to reach the oocyte boundary by stage 10. The migration of border cells requires integration of guidance cues with Rac1 GTPase activation, which leads to production of large migratory protrusions at the cluster leading edge (Fernández-Espartero et al., 2013; Montell et al., 2012; Ramel et al., 2013; Roberto and Emery, 2022; Saadin and Starz-Gaiano, 2016; Scarpa and Mayor, 2016; Wang et al., 2010). During the process of border cell collective migration, Rap1 promotes actomyosin polarity, helps restrict protrusions to the leading edge, and contributes to proper E-Cadherin enrichment within the cluster (Chang et al., 2018; Sawant et al., 2018). We know very little, however, about the Rap1 interacting genes that coordinate these critical Rap1-dependent functions.

Notably, similar to loss of Rap1 function, expression of constitutively active Rap1 (*Rap1^V12^*) causes severe border cell migration defects (Chang et al., 2018; Sawant et al., 2018). Thus, the levels of Rap1 are critical for normal collective cell migration. Here, we leveraged this phenotype to conduct an unbiased screen for genes whose heterozygous loss modifies the constitutively active Rap1 phenotype. Using a publicly available collection of *Drosophila* deficiencies that in total remove ∼98% of genes on the *X* chromosome, we identified seven genomic regions that dominantly suppressed the *Rap1^V12^* migration defects. Through further genetic tests, we mapped three of the interacting regions to individual genes. Specifically, we identified *frizzled 4 (fz4), Ubiquitin specific protease 16/45 (Usp16-45)*, and *strawberry notch (sno)* as genes whose heterozygous loss strongly modified the *Rap1^V12^*-induced border cell migration defects. Furthermore, we found that loss of *Usp16-45* and *sno* on their own also impaired border cell movement. Thus, this screen identified three genes and four additional genetic interacting regions that represent previously uncharacterized Rap1 interacting genes in collective cell migration.

## Methods & Materials

### *Drosophila* deficiency screen and genetics

The *X* chromosome deficiency kit (DK1) was obtained from the Bloomington *Drosophila* Stock Center (BDSC). Females from balanced *X* chromosome deficiency lines were crossed to *slbo-*GAL4/CyO; UAS-Rap1^V12^/TM6B, *tub*Gal80 males (Figure 1A). In the case of *w^1118^* controls, flies from the *slbo>*Rap1^V12^ stock were crossed to *w^1118^*. F1 progeny of the correct genotype (lacking balancer chromosomes) were selected and fattened overnight (∼12-24 hours) on supplemental yeast at 27°C prior to dissection. Each deficiency in the primary screen was tested at least once; potential interacting deficiency ‘hits’ were then further evaluated. Hits in the primary screen were defined as having at least 50% of border cell clusters migrating more than half of the egg chamber length (the region between the anterior tip and the oocyte anterior border; *see* Fig. 1A). This value is greater than two standard deviations above the mean of control egg chambers scored in the primary screen and allowed us to identify high confidence hits. All reported hit lines were subsequently tested a minimum of three times.

**Figure 1.**
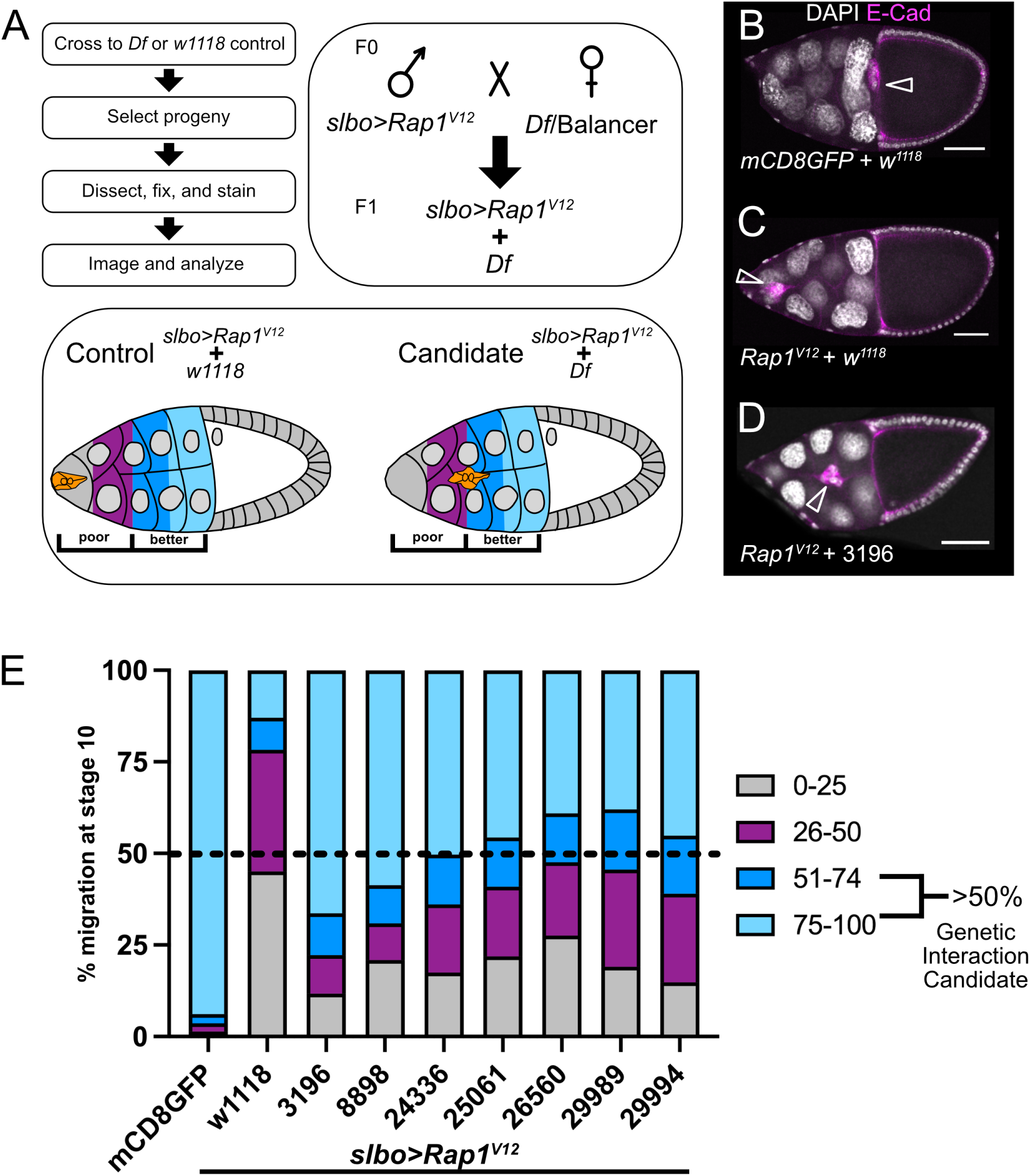
Screen to identify *Rap1^V12^* interacting regions. (A) Screen design flow, crossing scheme, and egg chamber schematic showing border cell migration scoring criteria. Poor migration includes border cells found from the anterior tip of the egg chamber to the midway point to the oocyte (gray, 0-25% migration distance to the oocyte; magenta, 26-50% migration distance to oocyte). Better migration includes border cells found from the midway point to the oocyte (dark blue, 51-74% migration distance to the oocyte; light blue, 75-100% migration distance to the oocyte). (B-D) Stage 10 egg chambers stained for E-cadherin (magenta) to label cell membranes and border cells (arrowheads) and DAPI to label cell nuclei (white). Scale bars, 50μm. (B) A representative *slbo>mCD8GFP* + *w^1118^* control egg chamber showing border cells (arrowhead) that completed their migration at the oocyte. (C) A representative *slbo>Rap1^V12^*+ *w^1118^* egg chamber that failed to migrate and stopped at ∼15% of the distance along the migration pathway. (D) An example of a candidate dominant modifier of Rap1^V12^. Here, *slbo>Rap1^V12^* + *Df(1)Sxl-bt* (3196) border cells exhibit better migration but stopped at the egg chamber midpoint. (E) Stacked bar chart displaying percent border cell migration at stage 10 for each of seven candidate deficiency regions in a *slbo*>*Rap1^V12^*background with *slbo>Rap1^V12^* + *w^1118^* and *slbo>mCD8GFP* + *w^1118^* serving as controls. N≥149 egg chambers per genotype. Colors represent the border cell migration distance to the oocyte as depicted in panel (A). The dashed line at 50% marks the screen hit threshold to be considered a genetic interaction candidate.

To identify relevant genes from the interacting deficiencies, UAS-RNAi lines and mutant alleles were obtained from BDSC and VDRC. For RNAi knockdown of candidate genes, each UAS-RNAi line was crossed to *c306-*GAL4, an early follicle cell driver that has been used for strong RNAi knockdown in border cells (Aranjuez et al., 2016; Miao et al., 2022; Plutoni et al., 2019). To ensure efficient RNAi knockdown, females of the correct genotype were temperature shifted to 29°C for two days before being fattened overnight (∼12-24 hours) on supplemental yeast at 29°C prior to dissection. Similarly, to test interaction with mutant alleles, *slbo-*GAL4/CyO; UAS-Rap1^V12^/TM6b, *tub*Gal80 males were crossed to mutant females, followed by incubation and fattening at 27°C prior to dissection and imaging of migration.

### Immunostaining and imaging

Ovaries were dissected in Schneider’s *Drosophila* Medium (Thermo Fisher Scientific, Waltham, MA, USA) supplemented with 10% fetal bovine serum (Seradigm FBS; VWR, Radnor, PA, USA). Ovaries were then fixed for 10 mins using 16% methanol-free formaldehyde (Polysciences, Inc., Warrington, PA, USA) diluted to a final concentration of 4% in 1X Phosphate Buffered Saline (PBS). Following fixation, tissues were washed ≥4x with ‘NP40 block’ (50 mM Tris-HCL, pH 7.4, 150mM NaCl, 0.5% NP40, 5mg/ml bovine serum albumin [BSA]) and rocked in the solution for ≥30 mins prior to antibody incubation. Primary antibodies, obtained from the Developmental Studies Hybridoma Bank (DSHB, University of Iowa, Iowa City, IA, USA), were used at the following dilutions: rat anti-E-Cadherin 1:10 (DCAD2) and mouse anti-Singed 1:10-1:25 (Sn7C). For GFP detection, rabbit anti-GFP (A11122, Thermo Fisher Scientific) was used at 1:1000 dilution. Anti-rat, anti-mouse, or anti-rabbit secondary antibodies conjugated to Alexa Flour-488 or -568 (Thermo Fisher Scientific) were used at 1:400 dilution. 4’, 6’-Diamidino-2-phenylindole (DAPI, Millipore Sigma) was used at 2.5 µg/ml to label nuclei. Primary and secondary antibody incubation as well as all other subsequent wash steps were also performed in NP40 block. Dissected and stained ovarioles and egg chambers were mounted on slides with Aqua-Poly/Mount (Polysciences, Inc.). Images of fixed egg chambers were either acquired on an upright Zeiss AxioImager Z1 microscope with Apotome.2 optical sectioning or on a Zeiss LSM 880 confocal laser scanning microscope (KSU College of Veterinary Medicine Confocal Core) using either a 20× 0.75 numerical aperture (NA) or 40× 1.3 NA oil-immersion objective controlled by Zeiss Zen 14 software. Images were processed in FIJI (Schindelin et al., 2012) and figures were assembled using Affinity Photo version 1.10.8 (Serif, Nottingham, United Kingdom). Illustrations were designed in Affinity Photo.

### Graphs and statistics

Deficiency regions that were considered “hits” were analyzed for migration a minimum of three times, with a minimum of 20 egg chambers scored per trial. The average of all trials for each “hit” had migration greater than 50% of the migration distance towards the oocyte. This value was determined as >2σ above the mean “migration” for *slbo*-GAL4, UAS-Rap1^V12^/*w^1118^* controls under the same conditions. To identify the relevant genes within the deficiency required for border cell migration in the RNAi tests, we defined a migration defect as the fraction of border cell clusters that failed to reach ≥75% of the migration distance to the oocyte. Chi-squared tests were performed to assess significance level for each experiment using GraphPad Prism 9 (GraphPad Software, San Diego, CA, USA). See Tables 2 and 3 for statistics. Graphs were assembled in GraphPad Prism 9. Relevant data from the screening approach were included in graphs where appropriate, but statistical testing was limited to experiments with matched controls.

**Table 1.** Primary screen results.

| Symbol | BDSC*<br>Stock<br># | Breakpoints | Est.<br>Cytolog<br>y | Hit<br>(Y/N)<br>^ | Fraction<br>Migrating<br>≤50% | Fraction<br>Migrating<br>>50% | #<br>Repe<br>ats | # EC§<br>scored |
| --- | --- | --- | --- | --- | --- | --- | --- | --- |
| Df(1)BSC843 | 27887 | X:254968-255277;X:33<br>4685 (Df) | 1A1;1A3<br>(Df) | N | 70.49% | 29.51% | 1 | 122 |
| Df(1)BSC530 | 25058 | X:364350;X:6<br>23478-623577 (Df) | 1A5;1B1<br>2 (Df) | N | 73.21% | 26.79% | 1 | 56 |
| Df(1)G1 | 34050 | X:644873;X:6<br>54238 (Df) | 1B13;1B<br>13 (Df) | N | 66.95% | 33.05% | 2 | 214 |
| Df(1)ED6443 | 9053 | X:656023;X:1<br>026707 (Df) | 1B14;1E<br>1 (Df) | N | 98.55% | 1.45% | 3 | 268 |
| Df(1)BSC534 | 25062 | X:841105;X:1<br>453730 (Df) | 1D1;2A<br>3 (Df) | N | 82.69% | 17.31% | 1 | 52 |
| Df(1)BSC719 | 26571 | X:1453730;X:<br>1865709 (Df) | 2A3;2B1<br>3 (Df) | N | 90.01% | 9.99% | 2 | 122 |
| Df(1)BSC717 | 26569 | X:2251580;X:<br>2545663 (Df) | 2F2;3A4<br>(Df) | N | 98.55% | 1.45% | 3 | 99 |
| Df(1)ED411 | 8031 | X:2469859;X:<br>2642686 (Df) | 3A3;3A8<br>(Df) | N | 70.63% | 29.38% | 1 | 160 |
| Df(1)ED6584 | 9348 | X:2636213;X:<br>2685435 (Df) | 3A8;3B1<br>(Df) | N | 97.26% | 2.74% | 3 | 199 |
| Df(1)ED6630 | 8948 | X:2685540;X:<br>3036910 (Df) | 3B1;3C<br>5 (Df) | N | 99.60% | 0.40% | 3 | 200 |
| Df(1)BSC531 | 25059 | X:2913683-2913782;X:3<br>672682 (Df) | 3C3;3E<br>2 (Df) | N | 52.83% | 47.17% | 1 | 53 |
| Df(1)BSC834 | 27886 | X:3288956;X:<br>3845727 (Df) | 3C11;3F<br>3 (Df) | N | 96.53% | 3.47% | 3 | 227 |
| Df(1)ED6712 | 9169 | X:3432535;X:<br>3789615 (Df) | 3D3;3F1<br>(Df) | N | 94.74% | 5.26% | 2 | 96 |
| Df(1)ED6716 | 24145 | X:3799196;X: | 3F2;4B3 | N | 94.54% | 5.46% | 3 | 242 |

| Symbol | BDSC*<br>Stock<br># | Breakpoints | Est.<br>Cytolog<br>y | Hit<br>(Y/N)<br>^ | Fraction<br>Migrating<br>≤50% | Fraction<br>Migrating<br>>50% | #<br>Repe<br>ats | # EC\$<br>scored |
| --- | --- | --- | --- | --- | --- | --- | --- | --- |
|  |  | 4204584 (Df) | (Df) |  |  |  |  |  |
| Df(1)BSC580 | 25414 | X:4101232-4101613;X:4<br>688080 (Df) | 4A5;4C<br>13 (Df) | N | 69.83% | 30.17% | 2 | 102 |
| Df(1)ED6727 | 8956 | X:4325174;X:<br>4911061 (Df) | 4B6;4D<br>5 (Df) | N | 85.71% | 14.29% | 1 | 49 |
| Df(1)JC70 | 944 | X:4679537-4919558;X:5<br>309242-5412386 (Df) | 4C12-4D6;4F4<br>-4F9<br>(Df) | N | 95.98% | 4.02% | 2 | 177 |
| Df(1)BSC533 | 25061 | X:5282581-5282584;X:5<br>428543 (Df) | 4F4;4F1<br>0 (Df) | Y | 40.92% | 59.08% | 4 | 333 |
| Df(1)Exel6235 | 7709 | X:5516611;X:<br>5593966 (Df) | 5A2;5A6<br>(Df) | N | 58.10% | 41.90% | 2 | 185 |
| Df(1)BSC571 | 25114 | X:5545559;X:<br>5662787 (Df) | 5A4;5A1<br>0 (Df) | N | 66.07% | 33.93% | 1 | 56 |
| Df(1)ED6802 | 8949 | X:5679980;X:<br>5965880 (Df) | 5A12;5<br>D1 (Df) | N | 56.07% | 43.93% | 2 | 172 |
| Df(1)ED6829 | 8947 | X:5901976;X:<br>6353095 (Df) | 5C7;5F3<br>(Df) | N | 77.50% | 22.50% | 1 | 80 |
| Df(1)Exel6239 | 7713 | X:6344333;X:<br>6516952-6538013 (Df) | 5F2;6B1<br>-6B2<br>(Df) | N | 51.06% | 48.94% | 2 | 127 |
| Df(1)Exel6240 | 7714 | X:6543963;X:<br>6669857-6669858 (Df) | 6B2;6C<br>4 (Df) | N | 60.13% | 39.87% | 1 | 153 |
| Df(1)BSC535 | 25063 | X:6625450;X:<br>6707019 (Df) | 6C2;6C<br>8 (Df) | N | 69.12% | 30.88% | 1 | 136 |
| Df(1)BSC351 | 24375 | X:6748387-6748403;X:6<br>860753 (Df) | 6C11;6<br>D7 (Df) | N | 97.36% | 2.64% | 3 | 189 |
| Df(1)BSC882 | 30587 | X:6824174;X: | 6D3;6E | N | 77.67% | 22.33% | 1 | 103 |

| Symbol | BDSC*<br>Stock<br># | Breakpoints | Est.<br>Cytolog<br>y | Hit<br>(Y/N)<br>^ | Fraction<br>Migrating<br>≤50% | Fraction<br>Migrating<br>>50% | #<br>Repe<br>ats | # EC<sup>s</sup><br>scored |
| --- | --- | --- | --- | --- | --- | --- | --- | --- |
|  |  | 7015408 (Df) | 4 (Df) |  |  |  |  |  |
| Df(1)BSC867 | 29990 | X:6981859;X:<br>7041515 (Df) | 6E4;6F1<br>(Df) | N | 85.68% | 14.32% | 2 | 188 |
| Df(1)Sxl-bt | 3196 | X:6987188-7004151;X:7<br>195487-7307939 (Df) | 6E4;7A3<br>-7B1<br>(Df) | Y | 22.20% | 77.80% | 5 | 439 |
| Df(1)ED6906 | 8955 | X:7195084;X:<br>7405806 (Df) | 7A3;7B2<br>(Df) | N | 73.82% | 26.18% | 2 | 96 |
| Df(1)BSC536 | 25064 | X:7338653;X:<br>7891613 (Df) | 7B2;7C<br>1 (Df) | N | 83.11% | 16.89% | 2 | 88 |
| Df(1)C128 | 949 | X:7901331-7956278;X:8<br>061645-8115848 (Df) | 7C2-7D1;7D<br>5-7D6<br>(Df) | N | 82.03% | 17.97% | 2 | 184 |
| Df(1)BSC866 | 29989 | X:8086993;X:<br>8157322 (Df) | 7D5;7D<br>16 (Df) | Y | 45.61% | 54.39% | 3 | 149 |
| Df(1)BSC662 | 26514 | X:8116248;X:<br>8489613 (Df) | 7D6;7F1<br>(Df) | N | 50.63% | 49.37% | 3 | 270 |
| Df(1)M38-C5 | 5706 | X:8877627-8933650;X:9<br>554623-9594143 (Df) | 8B5-8C1;8E<br>7-8E12<br>(Df) | N | 69.70% | 30.30% | 1 | 33 |
| Df(1)ED6957 | 8033 | X:8891795;X:<br>9135037 (Df) | 8B6;8C<br>13 (Df) | N | 75.63% | 24.37% | 1 | 119 |
| Df(1)BSC712 | 26564 | X:9606595;X:<br>10086569<br>(Df) | 8F1;9B1<br>(Df) | N | 76.02% | 23.98% | 3 | 186 |
| Df(1)ED7005 | 9153 | X:10071922;<br>X:10585431<br>(Df) | 9B1;9D<br>3 (Df) | N | 96.94% | 3.06% | 3 | 212 |
| Df(1)BSC755 | 26853 | X:10454979;<br>X:10848473 | 9C4;9F5<br>(Df) | N | 70.80% | 29.20% | 1 | 113 |
|  |  | (Df) |  |  |  |  |  |  |
| Df(1)BSC540 | 25068 | X:10772545;<br>X:11065010<br>(Df) | 9E8;10A<br>3 (Df) | N | 61.97% | 38.03% | 1 | 71 |
| Df(1)v-L1 | 6219 | X:10854869-10925631;X:<br>11108482-11136887<br>(Df) | 9F5-9F11;10<br>A4-10A6<br>(Df) | N | 74.60% | 25.40% | 1 | 63 |
| Df(1)BSC572 | 25391 | X:10890940;<br>X:11092253<br>(Df) | 9F8;10A<br>4 (Df) | N | 78.03% | 21.97% | 1 | 132 |
| Df(1)BSC287 | 23672 | X:11182121;<br>X:11426241<br>(Df) | 10A10;1<br>0B11<br>(Df) | N | 99.62% | 0.38% | 3 | 299 |
| Df(1)BSC722 | 26574 | X:11350466;<br>X:11754251<br>(Df) | 10B3;10<br>E1 (Df) | N | 93.00% | 7.00% | 2 | 89 |
| Df(1)ED7147 | 9171 | X:11714383;<br>X:12004800<br>(Df) | 10D6;11<br>A1 (Df) | N | 91.24% | 8.76% | 3 | 252 |
| Df(1)ED7161 | 9217 | X:12007087;<br>X:12750866<br>(Df) | 11A1;11<br>B14 (Df) | N | 93.11% | 6.89% | 3 | 247 |
| Df(1)ED7170 | 8898 | X:12752602;<br>X:13277326<br>(Df) | 11B15;1<br>1E8 (Df) | Y | 30.90% | 69.10% | 7 | 532 |
| Df(1)ED7225 | 24146 | X:13784406;<br>X:14322206<br>(Df) | 12C4;12<br>E8 (Df) | N | 100.00% | 0.00% | 1 | 61 |
| Df(1)ED7229 | 9352 | X:14222234;<br>X:14653944<br>(Df) | 12E5;12<br>F2 (Df) | N | 57.26% | 42.74% | 2 | 126 |
| Df(1)ED7261 | 9218 | X:14653809;<br>X:14839412<br>(Df) | 12F2;12<br>F5 (Df) | N | 87.67% | 12.33% | 5 | 424 |
| Df(1)BSC310 | 24336 | X:14842413;<br>X:15089556<br>(Df) | 12F5;13<br>A10 (Df) | Y | 36.07% | 63.93% | 4 | 175 |
| Df(1)ED7289 | 29732 | X:15024777;<br>X:15125750<br>(Df) | 13A5;13<br>A12 (Df) | N | 96.48% | 3.52% | 5 | 543 |
| Df(1)ED7294 | 8035 | X:15175415;<br>X:15450298<br>(Df) | 13B1;13<br>C3 (Df) | N | 69.33% | 30.67% | 1 | 150 |
| Df(1)ED7331 | 9219 | X:15450255;<br>X:15813523<br>(Df) | 13C3;13<br>F1 (Df) | N | 87.50% | 12.50% | 3 | 239 |
| Df(1)BSC714 | 26566 | X:15758351;<br>X:16086028<br>(Df) | 13E14;1<br>4A8 (Df) | N | 81.08% | 18.92% | 1 | 37 |
| Df(1)BSC758 | 26855 | X:16005260;<br>X:16367112<br>(Df) | 14A6;14<br>C1 (Df) | N | 53.67% | 46.33% | 2 | 191 |
| Df(1)BSC772 | 26869 | X:16302976;<br>X:16423105<br>(Df) | 14B9;14<br>C4 (Df) | N | 75.21% | 24.79% | 1 | 117 |
| Df(1)FDD-0024486 | 23295 | X:16423105;<br>X:16463156<br>(Df) | 14C4;14<br>D1 (Df) | N | 77.42% | 22.58% | 1 | 31 |
| Df(1)BSC760 | 26857 | X:16526332;<br>X:16632102<br>(Df) | 14E1;14<br>F2 (Df) | N | 50.14% | 49.86% | 2 | 154 |
| Df(1)BSC582 | 25416 | X:16680721;<br>X:17091833<br>(Df) | 15A1;15<br>E2 (Df) | N | 79.94% | 20.06% | 4 | 501 |
| Df(1)ED7374 | 8954 | X:16695187;<br>X:17107632<br>(Df) | 15A1;15<br>E3 (Df) | N | 91.33% | 8.67% | 3 | 253 |
| Df(1)BSC405 | 24429 | X:17830759-17830846;X:<br>18092832<br>(Df) | 16D5;16<br>F6 (Df) | N | 83.33% | 16.67% | 1 | 30 |
| Df(1)ED13478 | 29733 | X:18085406;<br>X:18102011<br>(Df) | 16F6;16<br>F7 (Df) | N | 88.55% | 11.45% | 2 | 218 |
| Df(1)BSC352 | 24376 | X:18117467;<br>X:18374885<br>(Df) | 16F7;17<br>A8 (Df) | N | 94.62% | 5.38% | 3 | 274 |
| Df(1)BSC716 | 26568 | X:18243732;<br>X:18800267<br>(Df) | 17A3;17<br>D6 (Df) | N | 84.64% | 15.36% | 2 | 148 |
| Df(1)ED7424 | 9350 | X:18657253;<br>X:19298773<br>(Df) | 17D1;18<br>C1 (Df) | N | 86.87% | 13.13% | 2 | 147 |
| Df(1)Exel7468 | 7768 | X:19264512;<br>X:19509637<br>(Df) | 18B7;18<br>C8 (Df) | N | 75.00% | 25.00% | 1 | 52 |
| Df(1)BSC275 | 23171 | X:19496689;<br>X:19580079<br>(Df) | 18C8;18<br>D3 (Df) | N | 100.00% | 0.00% | 1 | 22 |
| Df(1)BSC871 | 29994 | X:19617601;<br>X:19788713<br>(Df) | 18D7;18<br>F2 (Df) | Y | 38.96% | 61.04% | 4 | 193 |
| Df(1)BSC586 | 25420 | X:19788713;<br>X:20429928<br>(Df) | 18F2;19<br>D1 (Df) | N | 90.91% | 9.09% | 1 | 44 |
| Df(1)BSC644 | 25734 | X:20243402;<br>X:21061001 | 19C1;19<br>E7 (Df) | N | 80.49% | 19.51% | 1 | 41 |

| Symbol | BDSC*<br>Stock<br># | Breakpoints | Est.<br>Cytolog<br>y | Hit<br>(Y/N)<br>^ | Fraction<br>Migrating<br>≤50% | Fraction<br>Migrating<br>>50% | #<br>Repe<br>ats | # EC<sup>§</sup><br>scored |
| --- | --- | --- | --- | --- | --- | --- | --- | --- |
|  |  | (Df) |  |  |  |  |  |  |
| Df(1)DCB1-35b | 977 | X:20939503-21345088;X:<br>22980722-23542271<br>(Df) | 19E5-19F5;20<br>F3-h32<br>(Df) | N | 61.76% | 38.24% | 1 | 34 |
| Df(1)BSC708 | 26560 | X:21028296;<br>X:21623866<br>(Df) | 19E7;20<br>A4 (Df) | Y | 47.56% | 52.44% | 5 | 351 |
| Df(1)Exel6255 | 7723 | X:21519203;<br>X:22517665<br>(Df) | 20A1;20<br>C1 (Df) | N | 50.22% | 49.78% | 4 | 390 |

\*BDSC, Bloomington *Drosophila* Stock Center.
^Y, yes; N, no.
<sup>§</sup>EC, egg chambers.

**Table 2.** Candidate allele results.

| Allele | Sample size (N) | Migrating >50% | Statistical Test | Significance |
| --- | --- | --- | --- | --- |
| *fz4[3-1]* | 393 | 180 | Chi-square test | ****, P<0.0001 |
| matched control | 287 | 49 |  |  |
| *Usp16-45[B]* | 292 | 159 | Chi-square test | ****, P<0.0001 |
| matched control | 354 | 64 |  |  |
| *sno[EF531]* | 525 | 198 | Chi-square test | ****, P<0.0001 |
| matched control | 411 | 67 |  |  |
| *Su(H)[2]* | 410 | 173 | Chi-square test | ****, P<0.0001 |
| matched control | 287 | 49 |  |  |

N, number of egg chambers scored.

**Table 3.** Candidate gene RNAi results.

| RNAi Line/Gene | Sample size (N) | Migrating <75% | Statistical Test | Significance |
| --- | --- | --- | --- | --- |
| 64990 *fz4* RNAi #1 * | 238 | 0 | Chi-square test | **, P<0.01 |
| 60102 matched control^ | 202 | 9 |  |  |
| 102339 *fz4* RNAi #2^ | 186 | 9 | Chi-square test | ns |
| 60102 matched control^ | 239 | 9 |  |  |
| 41976 *Usp16-45* RNAi #1^ | 152 | 35 | Chi-square test | ****, P<0.0001 |
| 60102 matched control^ | 239 | 9 |  |  |
| 110286 *Usp16-45* RNAi #2^ | 196 | 4 | Chi-square test | ns |
| 60102 matched control^ | 239 | 9 |  |  |
| 28341 *sno* RNAi #1^ | 185 | 28 | Chi-square test | ****, P<0.0001 |
| 60102 matched control^ | 239 | 9 |  |  |
| 101404 *sno* RNAi #2^ | 109 | 39 | Chi-square test | ****, P<0.0001 |
| 60102 matched control^ | 259 | 9 |  |  |

N, number of egg chambers scored.
\*, BDSC stock
^, VDRC stock

## Results and Discussion

### A screen of the *X* chromosome for dominant modifiers of activated Rap1

Previous work demonstrated that regulated Rap1 activity is critical for its function in border cell migration and protrusion dynamics (Chang et al., 2018; Sawant et al., 2018). Expression of constitutively active Rap1 (*Rap1^V12^*) specifically blocks border cell migration, causes ectopic protrusions, and alters the enrichment of E-Cadherin and F-actin within the cluster. These previous results demonstrate a requirement for Rap1 in generating productive directed protrusions and promoting the distribution of cell-cell adhesions during border cell migration. Rap1 interacts with the Hippo/Warts pathway in directed protrusions (Chang et al., 2018). While other Rap1 effectors likely mediate additional Rap1-dependent functions in border cells, their identities are unknown. Thus, here we performed a dominant modifier screen to identify additional Rap1-interacting genes in border cell migration. The *X* chromosome was chosen on the basis that chromosomes *2* and *3* had been screened previously for modifiers of constitutively activated Rap1^V12^ (Chang et al., 2018).

For the primary screen we used the Bloomington Drosophila Stock Center (BDSC) *X* chromosome deficiency kit (DK1), which on aggregate deletes ∼98% of the euchromatic *X* chromosome (Cook et al., 2012). Of the 93 lines within this kit, 19 lines were unable to be tested due to complicated genetics and/or health issues of the stocks. We crossed the remaining female strains bearing the *X* chromosome deficiencies to males expressing UAS*-Rap1^V12^*driven by the border cell specific driver *slbo-GAL4* (“*slbo*>*Rap1^V12^*”; Figure 1A). Progeny expressing *Rap1^V12^* driven by *slbo-GAL4* outcrossed to *w^1118^* “wild type” background (“*slbo*>*Rap1^V12^* + *w^1118^*”) exhibited very strong border cell migration defects (Figure 1A, C, E). By contrast, *slbo-GAL4* driven expression of UAS-*mCD8-GFP* in a *w^1118^* background (“*slbo*>*mCD8GFP* + *w1118*”) resulted in normal migration (Figure 1B, E). To quantify the Rap1^V12^ migration defect severity, we divided each egg chamber into four quadrants, 0-25, 26-50, 51-74, and 75-100% of the normal migration distance from the anterior egg chamber tip (0%) to the oocyte (100%; Figure 1A, E). Relatively few *slbo*>*Rap1^V12^* + *w^1118^* border cell clusters traveled past the midpoint of migration (22% of egg chambers; Figure 1A, C, E). Therefore, we further simplified our scoring criteria by dividing the egg chamber into a region of “poor migration”, defined as 0-50% of the migration distance before the midpoint of migration, and a region of “better migration”, defined as 51-100% of the migration distance after the midpoint of migration (Figure 1A). Using these criteria, we then performed the screen for deficiencies on the *X* chromosome that modified the *slbo*>*Rap1^V12^*migration defects. We focused primarily on suppressors of *Rap1^V12^*because we reasoned that these genes were more likely to be downstream of Rap1. Therefore, we considered a screen hit as heterozygous loss of a deficiency that resulted in better migration in at least 50% of egg chambers assayed (Figure 1A, E).

Using this screening approach, we identified seven deficiency regions that dominantly suppressed the *slbo>Rap1^V12^* dependent migration defects (Figure 1A, D, E; Table 1). Each of these molecularly defined deficiencies completely or partially deletes an average of 9 genes, though some delete more genes (Cook et al., 2012). Using overlapping smaller deficiencies and publicly available mutant alleles and RNAi lines, we were able to map three of these regions to single interacting genes, which are described below.

### *Mapping the Df(1)Sxl-bt* region reveals an interaction between Rap1 and *fz4*

The strongest hit in this screen, *Df(1)Sxl-bt* (BDSC 3196), resulted in better migration in 78% of egg chambers analyzed (Figure 1 D, E; Table 1). This deficiency removes an estimated 191-321 kilobases (kb) of DNA (X:6987188-7004151 to X:7195487-7307939) along the *X* chromosome with a predicted deletion of 31 genes (FlyBase; Figure 2 A). We next used a smaller deficiency *Df(1)BSC867* (X:6,981,859 to X:7,041,515) to further refine the gene region (FlyBase; Figure 2 A). Border cell clusters expressing *slbo>Rap1^V12^*and heterozygous for *Df(1)BSC867* migrated past the midpoint only 14% of the time, similar to *slbo*>*Rap1^V12^* + *w^1118^*alone (Figure 2 C). We therefore considered it unlikely that Rap1 interacting genes resided within this segment of *Df(1)Sxl-bt*. We next focused on the region extending from the end of *Df(1)BSC867* to the end of *Df(1)Sxl-bt* (X:7,041,515-X:7,307,939). As there were no available deficiencies that overlapped with this region, we next tested for interaction with mutant alleles of characterized genes within the breakpoint region. Only two genes within this region, *Sex lethal* (*Sxl*) and *frizzled 4* (*fz4*) had characterized loss of function mutant alleles. Therefore, we tested interaction of *Sxl^f2^* and *fz4^3-1^* with Rap1^V12^. Border cells migrated past the midpoint in only 1% of *slbo>Rap1^V12^*+ *Sxl^f2^* egg chambers (Figure 2 C). In contrast, border cells in *slbo>Rap1^V12^* + *fz4^3-1^* egg chambers migrated significantly better, with 46% of clusters migrating past the midpoint compared to 18% in matched *slbo*>*Rap1^V12^* + *w^1118^* controls (Figure 2 B-C; Table 2; p<0.0001, Chi-squared test).

**Figure 2.**
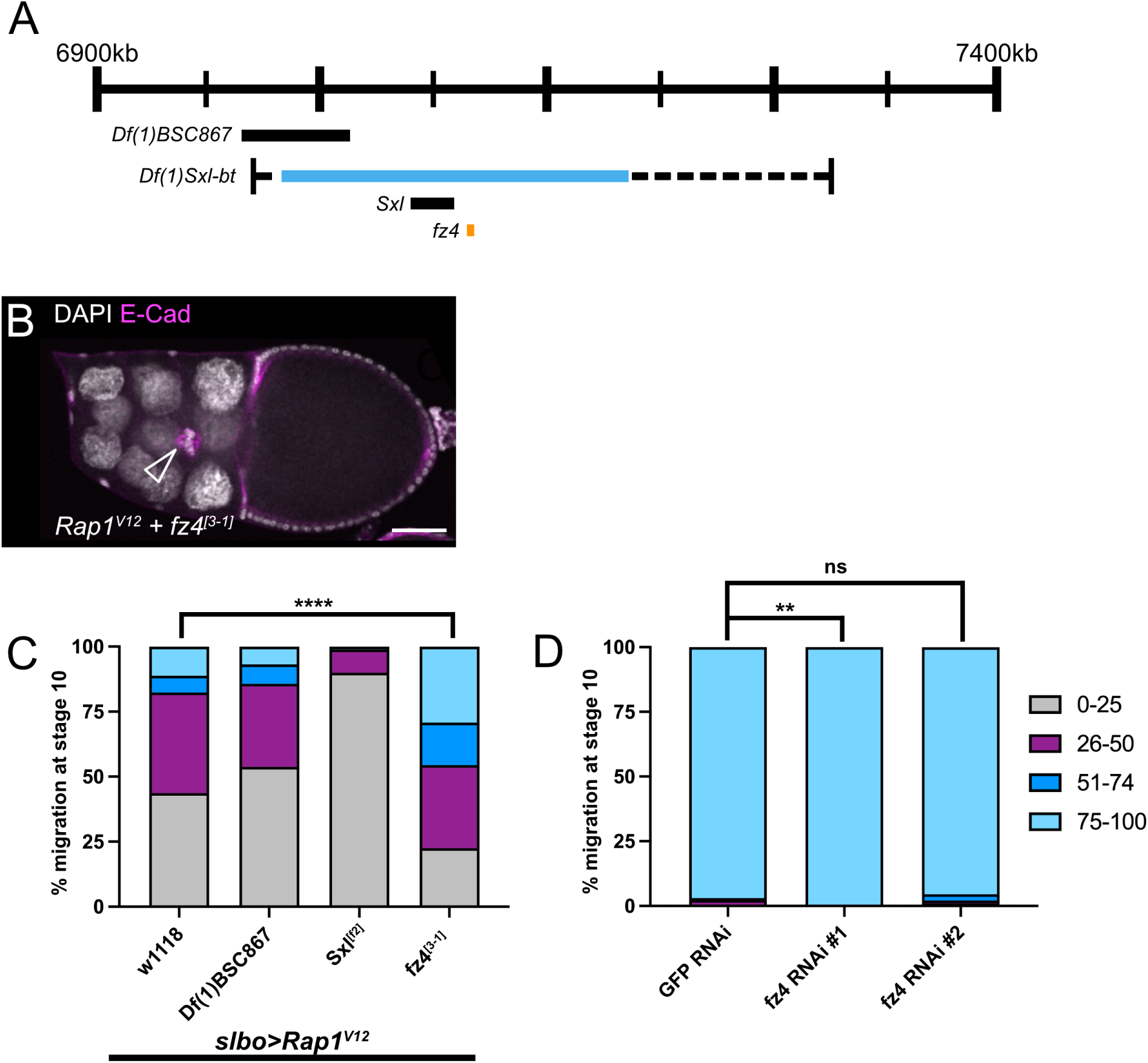
*Fz4* lies within *Df(1)Sxl-bt* and interacts with *Rap1^V12^*. (A) Schematic genomic region illustrating the location of *fz4* within *Df(1)Sxl-bt* along with overlapping deficiencies and genes tested. The numbers refer to the genomic location of the deficiency. (B) A stage 10 *Rap1^V12^* + *fz4^[3-1]^* egg chamber with border cells (arrowhead) that have moved past 50% of the egg chamber length. E-cadherin (magenta) labels all cell membranes including the border cells and DAPI labels cell nuclei (white). Scale bar, 50μm. (C) Stacked bar chart displaying border cell migration for *slbo>Rap1^V12^* + *w^1118^*, *slbo*>*Rap1^V12^* + *Df(1)BSC867* (BDSC 29990)*, slbo*>*Rap1^V12^* + *Sxl^[f2]^*(BDSC 4593), and *slbo*>*Rap1^V12^* + *fz4^[3-1]^*(BDSC 38412). N≥188 egg chambers per genotype. **** p<0.0001, two-sided Chi-square test. Data for *slbo*>*Rap1^V12^* + *Df(1)BSC867* (BDSC 29990) and *slbo*>*Rap1^V12^* + *Sxl^[f2]^*(BDSC 4593) are from the original mapping results and are shown here for simplicity. The statistical test was performed only between *slbo*>*Rap1^V12^* + *fz4^[3-1]^*(BDSC 38412) and the matched *slbo>Rap1^V12^* + *w^1118^* control (Table 2). (D) Stacked bar chart displaying border cell migration for *c306-GAL4>GFP* RNAi (VDRC 60102), *c306-GAL4*>*fz4* RNAi#1 (BDSC 64990), and *c306-GAL4*>*fz4* RNAi#2 (VDRC 102339). N≥186 egg chambers per genotype. ** p<0.001, two-sided Chi-square test; ns, not significant, two-sided Chi-square test. Cumulative *c306-GAL4>GFP* RNAi (VDRC 60102) data for this experiment is shown for simplicity, but statistical tests were performed between the matched experimental and control data (Table 3).

Single cell RNA sequencing data from the Fly Cell Atlas project revealed expression of *fz4* in both somatic and germline cells of the ovary, indicating that *fz4* is expressed in the relevant tissue (Li et al., 2022). Additionally, *fz4* transcript is differentially expressed during border cell migration (Burghardt et al., 2023). To determine whether *fz4* is required for border cell migration, we targeted *fz4* with RNAi using two independent, non-overlapping RNAi lines (*fz4* RNAi #1 BDSC 64990 and *fz4* RNAi #2 VDRC 102339), each of which were expressed under the control of a strong early follicle cell and border cell driver, *c306-GAL4* (Manseau et al., 1997)*. c306-GAL4* driven expression of a non-essential RNAi targeting GFP (“GFP RNAi”) resulted in normal migration, or ≥ 75% of the distance to the oocyte (Figure 2 D). Therefore, we considered any border cell clusters that failed to reach the oocyte (≤74% of the migration distance), as having a migration defect. *fz4* RNAi #1 resulted in normal migration, with all border cells migrating ≥75% of the distance to the oocyte (Figure 2 D). Similarly, *fz4* RNAi #2 resulted in a minimal 5% migration defect that resembled the 3% defect observed in GFP RNAi controls (Figure 2 D; Table 3). These data suggest that *fz4* on its own may be dispensable for the ability of border cells to complete their migration to the oocyte.

Fz4 is a member of the Frizzled family of proteins that act as receptors for secreted Wnt proteins (Huang and Klein, 2004). Wnt inhibitor of Dorsal (WntD/Wnt8) and Wnt4 both bind Fz4 (Gordon et al., 2005; McElwain et al., 2011; Wu and Nusse, 2002). WntD functions in the Toll-Dorsal pathway to pattern the gastrulating embryo but has limited expression in ovarian follicle cells (Ganguly et al., 2005; Li et al., 2022; Rahimi et al., 2016). Wnt4, however, contributes to cell movement in the pupal ovary and is required for border cell migration (Cohen et al., 2002; Kotian et al., 2022). The role for Wnt4 in border cell migration, however, may be independent of Fz4. RNAi for Fz4 did not impair migration, with the caveat that the knockdown efficiency may be incomplete. Alternatively, Wnt4 could bind multiple Frizzled proteins to coordinate its function in border cell migration. Indeed, in *Drosophila* S2 cells, Wnt4 binds to both Fz and Fz2 in addition to Fz4 (Wu and Nusse, 2002).

Currently, it is unclear how *fz4* heterozygosity modifies the *Rap1^V12^* border cell migration defect. Notably, *Rap1^V12^* border cell clusters accumulate excessive E-Cadherin at the cluster periphery (Sawant et al., 2018). Low levels of E-cadherin at the cluster periphery provide optimal traction of border cells upon the nurse cell substrate for forward movement (Cai et al., 2014; Niewiadomska et al., 1999). Wnt4 regulates Focal Adhesion Kinase (FAK) in the pupal ovary, a component of integrin-based focal adhesions (Chastney et al., 2024; Cohen et al., 2002). While Rap1 can regulate integrins in various cell types, FAK is not required for border cell migration and integrins appear to play minor roles in cluster organization (Grabbe et al., 2004; Llense and Martín-Blanco, 2008; Sun et al., 2022). It is possible that Wnt4, through Fz4 and possibly other Frizzled receptors, regulates multiple types of adhesions in migratory border cells (Kotian et al., 2022). Loss of *fz4*, therefore, could be sufficient to modify the adhesion defects caused by Rap1^V12^, but insufficient to cause border cell migration defects on its own.

### *Mapping the Df(1)BSC533* region reveals an interaction between Rap1 and *Usp16-45*

Border cell clusters expressing *slbo>Rap1^V12^* and heterozygous for *Df(1)BSC533* (BDSC 25061) migrated past the migration midpoint 59% of the time (Figure 1 E; Table 1). *Df(1)BSC533* deletes approximately 146 kb of DNA (X:5282581-5282584 to X:5428543) along the *X* chromosome and results in the predicted deletion of 21 genes (FlyBase; Figure 3 A). Two overlapping deficiencies, *Df(1)BSC823* (BDSC 27584; X:5,282,581 to X:5,332,808) and *Df(1)Exel6290* (BDSC 7753; X:5,364,532 to 5,428,543), each interacted with *slbo>Rap1^V12^* (FlyBase; Figure 3 A, D). While these two smaller deficiencies do not themselves overlap, the interaction with *Df(1)Exel6290* was stronger. Therefore, we focused on the *Df(1)Exel6290* region, which deletes six genes including *Ubiquitin specific protease 16/45 (Usp16-45;*Figure 3 A). A point mutation allele for this gene, *Usp16-45^[B]^*, was able to partially replicate the interaction observed for this deficiency with *Rap1^V12^*. Border cells migrated past the midpoint in 52% of *slbo>Rap1^V12^* + *Usp16-45^[B]^* egg chambers, compared to 19% in matched *slbo*>*Rap1^V12^* + *w^1118^* controls (Figure 3 B, D; Table 2; p<0.0001, Chi-squared test).

**Figure 3.**
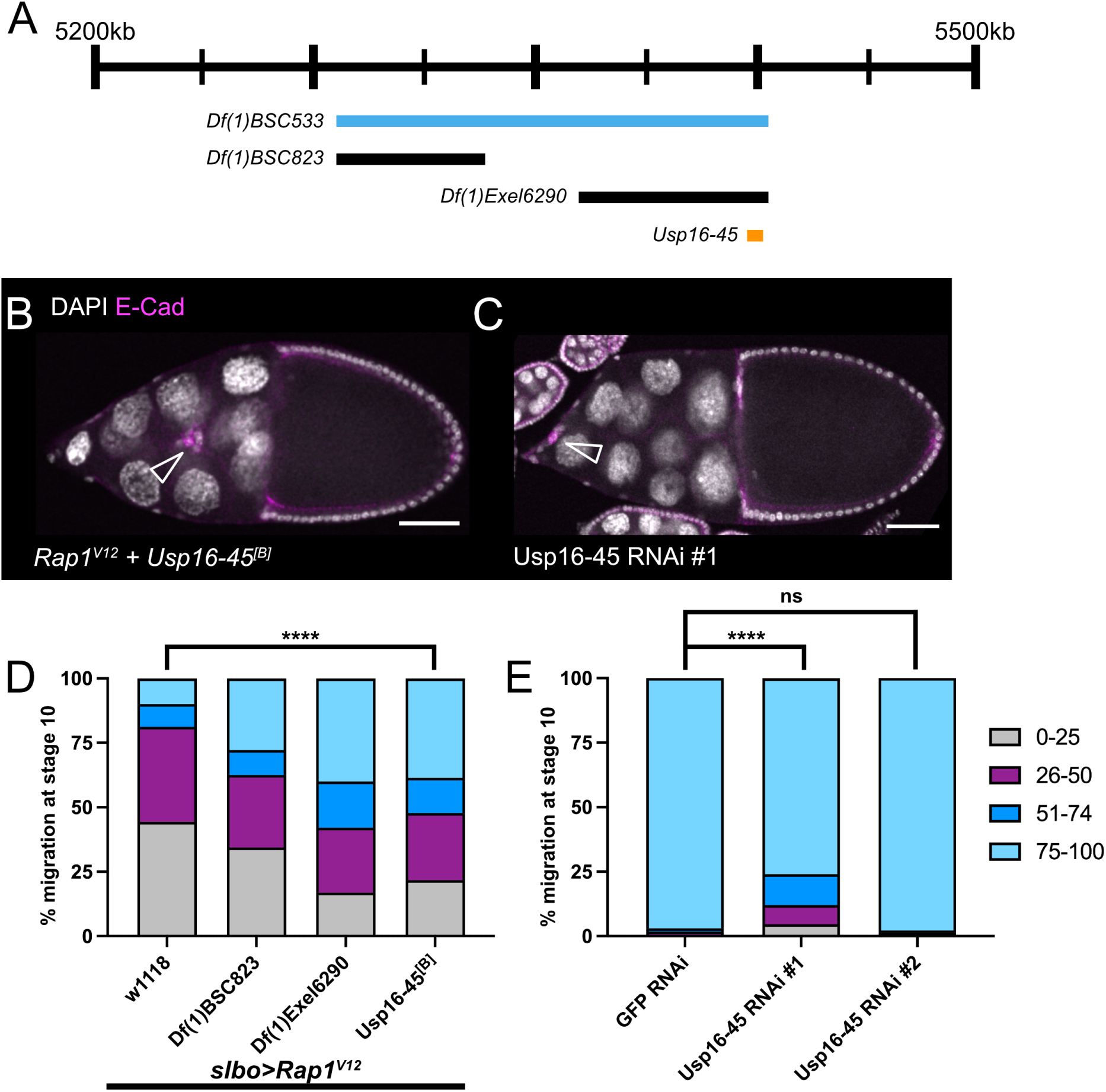
*Usp16-45* lies within *Df(1)BSC533,* interacts with *Rap1^V12^*, and is required for border cell migration. (A) Schematic genomic region illustrating where *Usp16-45* lies within *Df(1)BSC533* along with overlapping deficiencies tested. (B-C) Stage 10 egg chambers stained for E-cadherin (magenta), which labels all cell membranes including the border cells and DAPI to label cell nuclei (white). Arrowheads indicate border cell clusters. Scale bars, 50μm. (B) A *Rap1^V12^* + *Usp16-45^[B]^*egg chamber showing with border cells (arrowhead) moving past 50% of the egg chamber length. (C) A *c306-GAL4*>*Usp16-45* RNAi #1 (VDRC 41976) egg chamber with a strong border cell migration defect. (D) Stacked bar chart displaying border cell migration for *slbo>Rap1^V12^* + *w^1118^*, *slbo*>*Rap1^V12^* + *Df(1)BSC823* (BDSC 27584)*, slbo*>*Rap1^V12^* + *Df(1)Exel6290* (BDSC 7753), and *slbo*>*Rap1^V12^* + *Usp16-45^[B]^* (BDSC 57080). N≥182 egg chambers per genotype. **** p<0.0001, two-sided Chi-square test. Data for *slbo*>*Rap1^V12^* + *Df(1)BSC823* (BDSC 27584) and *slbo*>*Rap1^V12^* + *Df(1)Exel6290* (BDSC 7753) are from the original mapping approach and shown here for simplicity. The statistical test was performed only between *slbo*>*Rap1^V12^* + *Usp16-45^[B]^* (BDSC 57080) and *slbo>Rap1^V12^* + *w^1118^* matched control (Table 2). (E) Stacked bar chart displaying border cell migration for *c306-GAL4>GFP* RNAi (VDRC 60102), *c306-GAL4*>*Usp16-45* RNAi #1 (VDRC 41976), and *c306-GAL4*>*Usp16-45* RNAi #2 (VDRC 110286). N≥152 egg chambers per genotype. **** p<0.0001, two-sided Chi-square test; ns, not significant, two-sided Chi-square test (Table 3).

Usp16-45 is a member of the Ubiquitin Specific Proteases (USP) sub-family of deubiquitinases (Clague et al., 2019). While Usp16-45 has no known roles in cell migration, the family member non-stop (not), a USP22 ortholog, is required for border cell migration (Badmos et al., 2021). Single cell RNA sequencing data from the Fly Cell Atlas project revealed expression of *Usp16-45* in both somatic and germline cells of the ovary, indicating that *Usp16-45* is expressed in the relevant tissue (Li et al., 2022). *Usp16-45* is also differentially expressed in migrating border cells (Burghardt et al., 2023). To determine whether Usp16-45 is required for border cell migration, we expressed RNAi lines targeting the gene under control of *c306-*GAL4. Two independent, non-overlapping RNAi constructs (*Usp16-45* RNAi #1 VDRC 41976 and *Usp16-45* RNAi #2 VDRC 110286) provided mixed results. *Usp16-45* RNAi #1 resulted in moderately strong migration defects, with 24% of border cell clusters failing to reach the oocyte by stage 10 (Figure 3 C, E; Table 3). This value is significantly different than the 3% defect observed in controls (Figure 3 E; Table 3; p<0.0001, Chi-squared test). *Usp16-45* RNAi #2, however, resulted in minimal (2%) migration defects (Figure 3 E; Table 3). Although *Usp16-45* RNAi #2 failed to impact migration, it is possible that *Usp16-45* RNAi #1 results in more efficient knockdown of *Usp16-45.* Furthermore, no off-targets are predicted for *Usp16-45* RNAi #1, suggesting that the phenotypes observed are produced by specific knockdown of *Usp16-45* (Hu et al., 2013). Given the dominant genetic interaction of a *Usp16-45* mutant allele with Rap1^V12^ and the phenotypes caused by *Usp16-45* RNAi#1, we conclude that *Usp16-45* genetically interacts with Rap1 and is required for border cell migration.

Usp16-45 has predicted cysteine-type deubiquitinase activity (FlyBase; Komander et al., 2009). Small GTPases like Rap1 are often regulated by post-translational modifications (Konstantinopoulos et al., 2007). Post-translational modification at the CAAX domain, for example, can facilitate membrane targeting of GTPases (Konstantinopoulos et al., 2007). Ubiquitination is a mode of post-translational modification that can regulate small GTPase stability, activity, and localization (Lei et al., 2021). However, it is currently unclear whether Usp16-45 directly targets Rap1. RAPGEF2, an activating GEF for Rap1, is known to be targeted for ubiquitination (Kim et al., 2015). The RAPGEF2 ortholog PDZ-GEF (also known as Dizzy) is required for border cell migration, though a role for ubiquitination has not been tested (Sawant et al., 2018). These data suggest that the addition or removal of ubiquitin could be critical in regulating Rap1 signaling in border cells. Further work will be required to determine if Usp16-45 targets Rap1 directly, or indirectly via regulation of a signaling partner such as PDZ-GEF or another Rap1-interacting protein.

### *Mapping the Df(1)ED7170* region reveals an interaction between Rap1 and *sno*

Border cell clusters expressing *slbo>Rap1^V12^* and heterozygous for *Df(1)ED7170* (BDSC 8898) migrated past the midpoint in 69% of egg chambers (Figure 1 E; Table 1). *Df(1)ED7170* deletes approximately 525 kb of DNA (X:12,752,602 to X:13,277,326) along the *X* chromosome (FlyBase; Figure 4 A) and interacted strongly with *slbo>Rap1^V12^*. *Df(1)ED7170* is predicted to delete or disrupt 60 genes. Using deficiencies *Df(1)ED7165* (BDSC 9058; X:12,752,602 to X:13,138,948) and *Df(1)BSC713* (BDSC 26565; X:13,159,870 to X:13,373,704) led us to focus on the region extending from X:13,159,870 to X:13,277,326, which covers the beginning of *Df(1)BSC713* to the end of *Df(1)ED7170* (FlyBase; Figure 4 A, H). Using available alleles of genes within this region led us to investigate the gene *strawberry notch (sno)*. Sno is a nuclear protein that functions in Notch signaling (Majumdar et al., 1997). Single cell RNA sequencing data from the Fly Cell Atlas project indicates *sno* expression in both the germline and somatic cells of the ovary (Li et al., 2022). Using a GFP protein trap in *sno*, *sno*^CC01032^, we found that Sno is found in the nuclei of all cells including the nurse cells, follicle cells, and border cells (Figure 4 B, B’).

**Figure 4.**
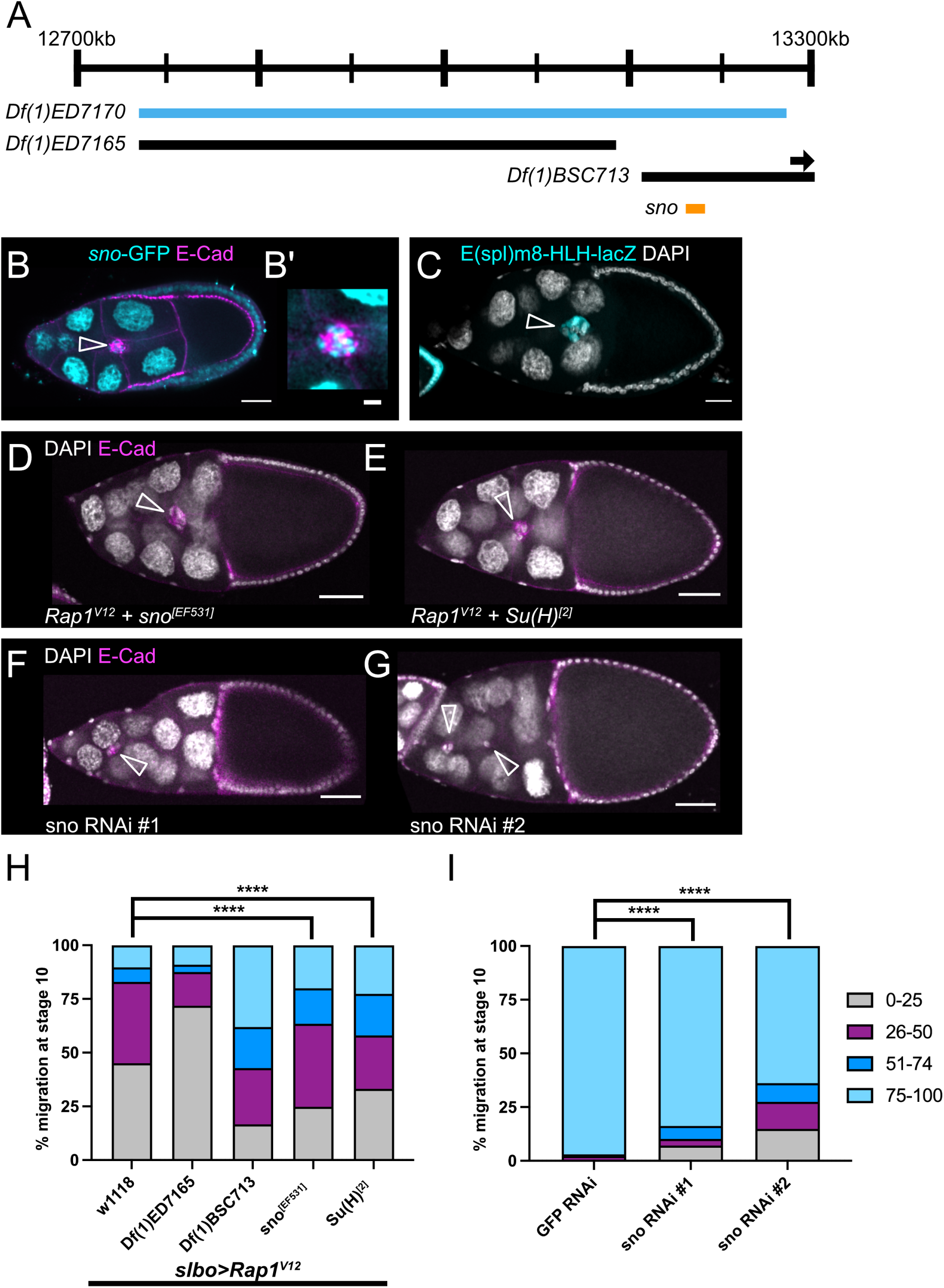
*Sno* lies within *Df(1)ED7170,* interacts with *Rap1^V12^*, and is required for border cell migration. (A) Schematic genomic region illustrating where *sno* lies within *Df(1)ED7170* along with overlapping deficiencies tested. Arrow indicates that *Df(1)BSC713* extends beyond the genomic region depicted here. (B) Egg chamber with GFP protein trap in *sno*, *sno*^CC01032^, shows Sno nuclear expression (cyan). E-cadherin (magenta) labels cell membranes including the border cells. (B’) Close-up view of the same border cell cluster in B. (C) Egg chamber with Notch activity reporter, *E(spl)m8-HLH-lacZ* (also known as Su(H)-lacZ), which shows nuclear expression in border cells (cyan). DAPI (white) labels cell nuclei. (D-G) Stage 10 egg chambers stained for E-cadherin (magenta), which labels all cell membranes including the border cells and DAPI to label cell nuclei (white). (D) A *Rap1^V12^* + *sno^[EF531]^*egg chamber with the border cells moving past 50% of the egg chamber length. (E) A *Rap1^V12^* + *Su(H)^[2]^* egg chamber with the border cells moving past 50% of the egg chamber length. (F) A *c306-GAL4*>*Sno* RNAi #1 (VDRC 28341) egg chamber with a strong border cell migration defect. (G) A *c306-GAL4*>*Sno* RNAi #2 (VDRC 101404) egg chamber with a strong border cell migration defect. (B-G) Arrowheads indicate border cell clusters. Scale bars, 20 µm (B and C), 5µm (B’), and 50μm (D-G). (H) Stacked bar chart displaying border cell migration for *slbo>Rap1^V12^*+ *w^1118^*, *slbo*>*Rap1^V12^* + *Df(1)ED7165* (BDSC 9058)*, slbo*>*Rap1^V12^* + *Df(1)BSC713* (BDSC 26565)*, slbo*>*Rap1^V12^* + *sno^[EF531]^* (BDSC 33833)*, slbo*>*Rap1^V12^* + *Su(H)^[2]^* (BDSC 30477). N≥42 egg chambers per genotype. **** p<0.0001, two-sided Chi-square test. Data for *slbo*>*Rap1^V12^* + *Df(1)ED7165* (BDSC 9058) and *slbo*>*Rap1^V12^* + *Df(1)BSC713* (BDSC 26565) are from the original mapping approach and shown here for simplicity. Statistical tests were performed only between *slbo*>*Rap1^V12^* + *sno^[EF531]^* (BDSC 33833) and *slbo>Rap1^V12^* + *w^1118^* matched control or *slbo*>*Rap1^V12^* + *Su(H)^[2]^* (BDSC 30477) and *slbo>Rap1^V12^* + *w^1118^* matched control (Table 2). (I) Stacked bar chart displaying border cell migration for *c306-GAL4>GFP* RNAi (VDRC 60102), *c306-GAL4*>*sno* RNAi #1 (VDRC 28341), and *c306-GAL4*>*sno* RNAi #2 (VDRC 101404). N≥109 egg chambers per genotype. **** p<0.0001, two-sided Chi-square test. Cumulative *c306-GAL4>GFP* RNAi (VDRC 60102) data for this experiment was reported for simplicity, but statistical tests were performed between matched experimental and control data (Table 3).

We were able to partially replicate the interaction observed for *Df(1)ED7170* with a loss of function allele for *sno, sno^EF531^*. Border cells migrated past the midpoint in 37% of egg chambers scored for *slbo>Rap1^V12^* + *sno^EF531^* compared to 17% in matched controls (Figure 4 D, H; Table 2; p<0.0001, Chi-squared test). Genetic interaction experiments in the wing and eye place *sno* in the Notch pathway (Coyle-Thompson and Banerjee, 1993). Rough eye and wing notching phenotypes found in *sno* mutants are rescued by an extra copy of *Notch* (Coyle-Thompson and Banerjee, 1993). Similarly, combining the hypomorphic *nd^1^*allele of *Notch* with the temperature sensitive *sno^71e3^*allele synergistically enhances the mild wing phenotypes present in *nd^1^*alone (Coyle-Thompson and Banerjee, 1993). Sno also binds to Suppressor of Hairless Su(H) downstream of Epidermal Growth Factor Receptor (EGFR) signaling in *Drosophila* eye development (Tsuda et al., 2002). Notch activity, as visualized by a lacZ reporter containing both Su(H) and Grainyhead (Grh) binding sites, E(spl)m8-HLH-lacZ, is high in migrating border cells (Figure 4C; Furriols and Bray, 2001; Schober et al., 2005; Wang et al., 2007). Therefore, to determine if the interaction between *sno* and *Rap1^V12^* is related to Sno-dependent regulation of Su(H), we next tested a *Su(H)* loss of function allele, *Su(H)^[2]^.* Border cells migrated past the midpoint in 42% of *slbo>Rap1^V12^* + *Su(H)^[2]^*egg chambers compared to 17% in *slbo*>*Rap1^V12^* + *w^1118^*controls (Figure 4 E, H; Table 2). These results indicate that *sno* and *Su(H)* both interact with Rap1 in border cell migration, possibly through their roles in the Notch pathway.

We next asked whether *sno* was essential for border cell migration on its own. Prior work indicates a role for Sno in successful oogenesis. Females homozygous for the *sno* allele *sno^71e1^*had severe defects in oogenesis including a reduced number of ovarioles, dying egg chambers, and disrupted egg chamber polarity (Coyle-Thompson and Banerjee, 1993). To determine whether *sno* is required specifically for border cell migration, we targeted *sno* with RNAi lines expressed under the control of the follicle cell driver *c306-GAL4*. Using two independent, non-overlapping constructs (*sno* RNAi #1 VDRC 28341 and *sno* RNAi #2 VDRC 101404), we found that *sno* is required for migration (Figure 4 F, G, I). *sno* RNAi #1 resulted in a significant migration defect of 16% (Figure 4 F, I; Table 3). Similarly, we observed 36% migration defect for *sno* RNAi #2 (Figure 4 G, I; Table 3). The difference in migration defects between RNAi lines is likely due to differences in knockdown efficiency.

How *sno* and *Su(H)* contribute to border cell migration via the Rap1 pathway is unclear. Notch and its ligand Delta are required for normal border cell migration (Schober et al., 2005; Wang et al., 2007). Both active Notch and *Su(H)* are expressed during migration (Figure 4C; Wang et al., 2007). One downstream target of Notch-Su(H) is Anterior open (Aop, also known as Yan; Schober et al., 2005). Aop regulates the turnover of E-Cadherin required for efficient border cell migration (Schober et al., 2005). One possibility is that Sno could regulate Aop, which in turn impacts migration efficiency via E-Cadherin turnover. The interaction between *sno* and *Rap1^V12^* could thus be explained by a common target, E-Cadherin. Further work will be required to determine if this hypothesis is supported, or if another mechanism is at play.

### Conclusions

The goal of this screen was to identify molecular partners and potential effectors of Rap1 GTPase activity that are relevant for collective cell migration. Here we report seven deficiency regions that genetically interacted with *slbo>Rap1^V12^*. Of these seven regions, three were mapped to single genes. Fz4, Usp16-45, and Sno may function as effectors of Rap1 GTPase or act as parallel factors that similarly regulate border cell migration. *Sno* and *Usp16-45* were each required for border cell migration on their own, while *fz4* was not. It is important to note that heterozygous loss of these genes did not completely recapitulate the interaction of the relevant deficiency with Rap1^V12^. This could be due to the heterozygous loss of additional genes in these genomic regions that interact with Rap1 or because of the nature of the mutant alleles used, though additional work is needed to determine which of these two possibilities is most likely. Future identification of the relevant genes from the four other interacting deficiencies is also expected to yield additional Rap1 effectors in border cells. In sum, this unbiased genetic modifier screen identified multiple genes that interact with Rap1 during border cell migration. Follow up studies will be needed to fully characterize how each gene cooperates with Rap1 to facilitate border cell migration and determine if these genes function in other types of collective cell migration during development and in cancer.

## Data Availability Statement

*Drosophila* strains are available upon request. The authors affirm that all data necessary for confirming the conclusions of the article are present within the article, figures, and tables. Table 1 contains the complete results of the screen. Table 2 shows the results of genetic interaction with mutant alleles and Table 3 shows results of RNAi knockdown in border cells.

## Acknowledgments

We would like to thank the Bloomington Drosophila Stock Center and the Vienna Drosophila Resource Center for providing flies, and the Developmental Studies Hybridoma Bank at the University of Iowa for providing antibodies used in this study. We used FlyBase (release FB2024_03) for information on genes, functions, and stocks. We also thank the Kansas State University Confocal Core for use of the Zeiss LSM880 confocal.

